# Dissecting the properties of circulating IgG against Group A Streptococcus through a combined systems antigenomics-serology workflow

**DOI:** 10.1101/2023.11.07.565977

**Authors:** Sounak Chowdhury, Alejandro Gomez Toledo, Elisabeth Hjortswang, James T Sorrentino, Nathan E Lewis, Anna Bläckberg, Simon Ekström, Arman Izadi, Pontus Nordenfelt, Lars Malmström, Magnus Rasmussen, Johan Malmström

**Author notes:** Corresponding author: Johan Malmström. Equal contribution (Sounak Chowdhury, Alejandro Gomez Toledo).

## Abstract

Most individuals maintain circulating antibodies against various pathogenic bacteria as a consequence of previous exposures. However, it remains unclear to what extent these antibodies contribute to host protection. This knowledge gap is linked to the need for better methods to characterize antimicrobial polyclonal antibodies, including their antigen and epitope repertoires, subclass distribution, glycosylation status, and effector functions. Here, we showcase a generic mass spectrometry-based strategy that couples systems antigenomics and systems serology to characterize human antibodies directly in clinical samples. The method is based on automated affinity purification workflows coupled to an integrated suite of high-resolution MS-based quantitative, structural- and glyco-proteomics readouts.

We focused on *Streptococcus pyogenes* (Group A *Streptococcus*; GAS), a major human pathogen still awaiting an approved vaccine. Our methodology reveals that both healthy and GAS infected individuals have circulating Immunoglobulin G (IgG) against a subset of genomically conserved streptococcal proteins, including numerous toxins and virulence factors. The antigen repertoire targeted by these antibodies was relatively constant across healthy individuals, but considerably changed in GAS bacteremia. Detailed analysis of the antigen-specific IgG indicates inter-individual variation regarding titers, subclass distributions, and Fc-signaling capacity, but not in epitope and Fc-glycosylation patterns. Importantly, we show that the IgG subclass has a major impact on the ability of GAS-antibodies to trigger immune signaling, in an antigen- and Fc receptor-specific fashion. Overall, these results uncover exceeding complexity in the properties of GAS-specific IgG, and showcase our methodology as high-throughput and flexible workflow to understand adaptive immune responses to bacterial pathogens.

**Significance statement:** Most people develop polyclonal antibodies against bacterial pathogens during infections but their structural and functional properties are poorly understood. Here, we showcase a combined systems antigenomics and systems serology strategy to quantify key antibody properties directly in clinical samples. We applied this method to characterize polyclonal antibody responses against S*treptococcus pyogenes*, a major human pathogen. We mapped the antigen and epitope landscape of anti-streptococcal antibodies circulating in healthy adult plasma, and their changes during blood infections. We further demonstrate the analytical power of our approach to resolve individual variations in the structure and effector functions of antigen-specific antibodies, including a dependency between immunoglobulin subclass and Fc-signaling capacity.

## Introduction

Immunoglobulin G (IgG) is a central effector molecule of adaptive immunity that leverages protective responses against microbial infections. IgG binds to the surface of viral and bacterial pathogens, and to soluble toxins, to neutralize their capacity to damage host tissues. Neutralization is mediated by the fragment antigen-binding (Fab) region, which recognizes epitopes on microbial proteins and polysaccharides. Neutralizing Fab binding prevents key steps in the establishment of an infection, including pathogen adhesion and cellular invasion. Besides neutralization, antigen-bound IgG can also trigger the initiation of the classical complement pathway, as well as other protective responses, such as antibody-dependent cellular cytotoxicity (ADCC) and antibody-dependent cellular phagocytosis (ADCP)(1). These effector functions are finetuned by the structure of the fragment crystallizable (Fc) region, especially by the Fc subclass and glycosylation, which synergistically modulate the IgG affinity for complement and immune cell receptors(2).

During antimicrobial responses, polyclonal IgG targets several antigens on a given pathogen, and various epitopes within each antigen, resulting in a broader range of protective responses compared to monoclonal IgG. However, characterizing the properties of polyclonal antibodies at a systems-wide level, including their antigenic repertoires, binding epitopes, subclass distributions, glycosylation patterns, and effector functions, remains a significant analytical challenge(3). In turn, a poor understanding of the structural and functional IgG features that contribute to host protection prevents the identification of useful correlates of immunity to major human pathogens, and the development of antimicrobial vaccines(4, 5).

Recently, efforts in reverse vaccinology have led to the development of systems antigenomic approaches that exploit the availability of annotated genome data, novel surface display technologies, and proteomics workflows, to characterize microbial antigens recognized by antibodies and T-cells (6–11). Systems antigenomics has been successful in defining pathogen-specific antibody antigenomes, (i.e., the spectrum of molecules expressed by a given pathogen that are recognized by host antibodies), a central bottleneck of most vaccine development pipelines(12, 13). However, with an obvious focus on antigen identification, systems antigenomics does not inform on other antibody properties beyond Fab binding. Advances in Omics technologies have also sparked the field of systems serology, a collection of integrative approaches to analyze various antibody features and functions, coupled to advanced computational and statistical methods(14–17). Systems serology has been useful to deconvolute immune correlates of protection and vaccine efficacy for the Human Immunodeficiency Virus (HIV)(18), *Mycobacterium tuberculosis* (MTB)(19), and SARS-CoV-2(20, 21). However, the starting point of systems serology is typically one or a few preselected antigen(s), a choice that often relies on previously acquired knowledge.

Mass spectrometry (MS) is a highly sensitive and flexible analytical method to identify proteins, to measure protein abundances, and to characterize both post-translational modifications and protein-protein interactions. We hypothesized that the wide flexibility of modern MS technologies opens new opportunities to combine systems antigenomics and systems serology, to provide an unbiased way to identify relevant antigens, followed by a focused multilayered characterization of the antigen-specific antibodies. Here, we demonstrate an automated and quantitative workflow based on combined principles from both systems antigenomics and systems serology, and applied to analyze antibody responses against Group A *Streptococcus* (GAS), a major bacterial pathogen and a significant source of human morbidity and mortality worldwide(22).

## Results

Most adult individuals have circulating IgG antibodies against GAS, but their structural and functional properties remain poorly understood(23). To address this challenge, we developed a two-step approach to i) determine the GAS-antigenome, and ii) characterize the titers, epitope repertoires, immune signaling capacity, subclass distributions, and N-linked glycosylation profiles of the antigen-specific IgG **(Fig. 1A).** This approach was built on streamlining antigen/antibody affinity purification workflows using an automated liquid-handling platform, coupled to a suite of high-resolution MS-based quantitative, structural- and glyco-proteomics readouts.

**Fig.1.**
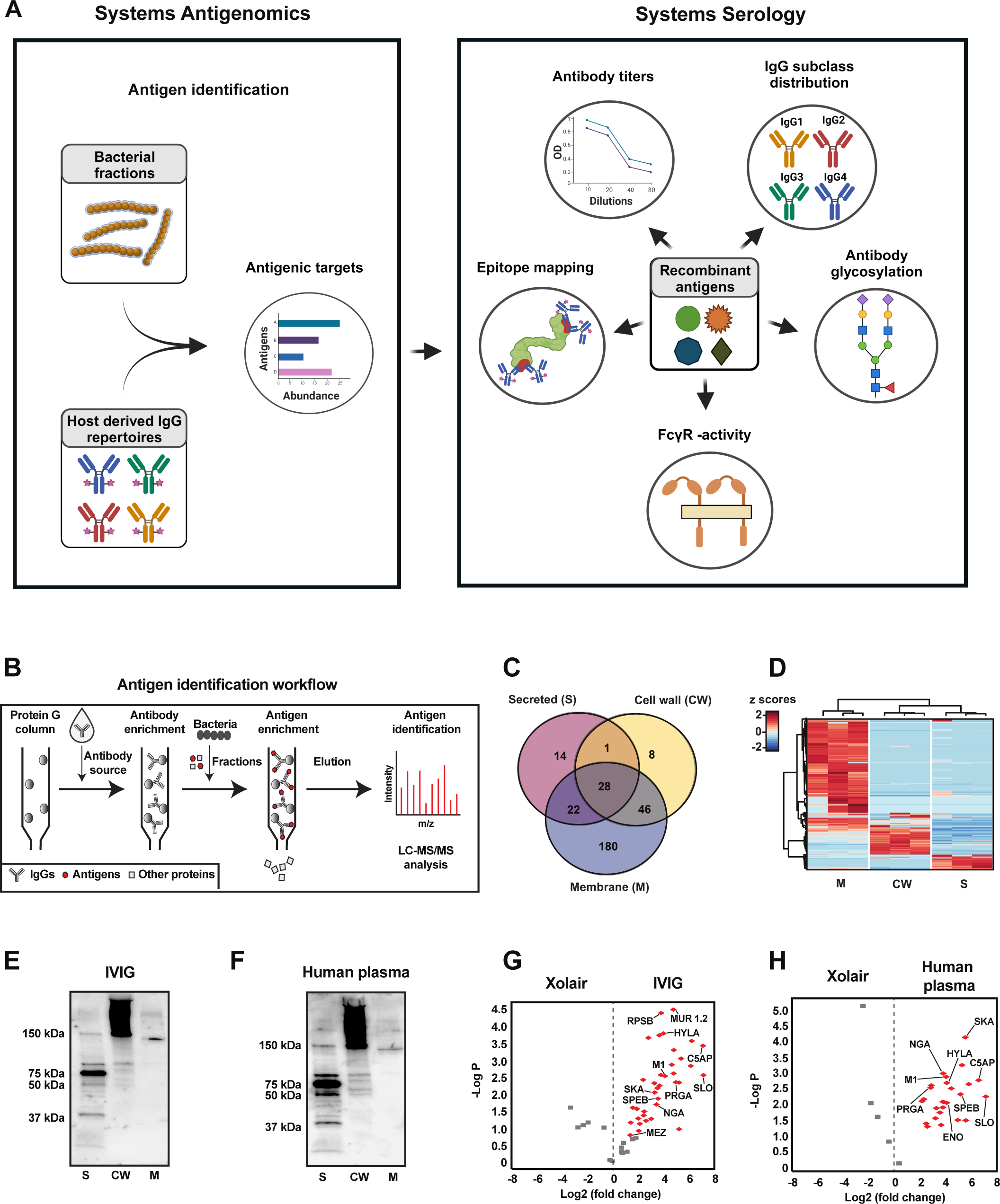
The GAS-specific IgG antigenome. **(A)** Schematic representation of the two-step approach integrating systems antigenomics and systems serology. The systems antigenomics strategy involves the identification of antigenic targets from biochemical fractions of bacterial proteins using host derived IgG as a guide. Selected antigens are then recombinantly produced and analyzed in a streamlined workflow of various systems serology techniques to deconvolute structural and functional attributes of the antigen specific IgG. **(B)** Schematic summary of the antigen identification workflow used in this study. **(C)** Overlap of the bacterial proteins identified across secreted (S), cell wall (CW) and membrane (M) fractions during a typical biochemical fractionation of the SF5370 GAS proteome. **(D)** Differential protein expression across bacterial fractions. The protein values were normalized using a Z-score normalization and subjected to Pearson correlation clustering. **(E)** Immunoblot analysis of GAS antigens in S, CW and M fractions using IVIG. **(F)** Immunoblot analysis of GAS antigens in S, CW and M fractions using pooled human plasma. Immunoblots are representative images of at least 2 independent experiments. **(G)** Volcano plot displaying the significant antigens recognized by IVIG and **(H)** pooled human plasma. Statistically significant identifications were assessed by 2-way ANOVA with a Bonferroni correction for multiple testing.

### Mapping the GAS- antigenome

To define the GAS-antigenome, we exploited GAS-specific IgG circulating in human plasma as a tool to isolate antigens from pools of bacterial proteins via affinity purification coupled to LC-MS/MS (**Fig. 1B**). First, the SF370 strain, a clinically relevant M1 GAS serotype, was biochemically fractionated into defined pools of potentially antigenic proteins **(Fig. S1A-E)**. The identity and cellular localization of bacterial proteins in pools from a typical preparation are presented in **Fig. 1C-D and supplemental table 1.** We primarily focused on surface exposed and secreted proteins since they are more likely to be recognized by host antibodies. Next, we used two IgG sources to isolate antigens from these bacterial fractions: i) pharmaceutical-grade intravenous immunoglobulin G (IVIG), and ii) commercial pooled human plasma (HP). Both IVIG and HP contain IgG antibodies from many different individuals, including clones directed against multiple GAS antigens **(Fig. 1E-F)**. IgG was immobilized on Protein G columns, and the bacterial fractions were passed through the columns to enrich for GAS antigens. Retained antigens were eluted and characterized by LC-MS/MS. This antigenomics approach identified a total of 39 antigens: 13 were unique to the IVIG, 2 were unique to HP, and 24 were common to both IVIG and HP **(Fig. 1G-H & supplemental table 2)**. The method was highly reproducible and generated a similar number of identifications across replicates (**Fig. S2A-D**). The GAS antigenome was enriched in virulence factors, including toxins (e.g., SLO), superantigens (e.g., SPEC, MEZ), anti-phagocytic proteins (e.g., M1), and enzymes (e.g., C5AP, HYLA). A few proteins of unknown functions (e.g., PRGA) or without an obvious link to GAS virulence (e.g., ribosomal RPLA and RPSB) were also identified (**Supplemental table 2)**. We conclude that GAS-specific IgG circulating in human plasma targets a small subset of bacterial antigens, many of which are well-known virulence determinants.

### The GAS-antigenome is conserved across healthy individuals but is altered in GAS bacteremia

In the next step, we analyzed plasma from 10 healthy donors to investigate potential individual variations in the GAS antigenome. In addition, both acute and convalescent plasma from four patients with GAS bacteremia were included to determine whether invasive infections might affect the antigenome profiles. We have previously reported the clinical and serological status of these patients, and no major differences were found between their acute and convalescent plasma(24). The antigenome analysis resulted in the identification of a total of 72 antigens, with an average of ∼30 antigens/individual. A substantial overlap was observed across the individual antigenomes, as well as between the individual and the “pooled” (i.e., IVIG and HP) antigenomes. **(Supplemental table 3).** The level of antigen enrichment varied across samples and correlated with antibody titers measured by ELISA, as evaluated for two antigens: C5AP and PRGA (**Fig. S3**). The antigenome profiles were rank correlated and linked by network analysis using Kendall Tau coefficients (**Fig. 2A**). The networking approach segregated the data into two distinct clusters. Cluster A was driven by antigens enriched in the patients with GAS bacteremia (e.g., ENO, LDH) **(Fig. 2B),** whereas Cluster B was driven by antigens enriched in the healthy group (e.g., MF, PRSA1) **(Fig. 2C)**. However, independently of disease status, all individuals had circulating antibodies against a common set of 11 antigens, suggesting a potential immunological signature of GAS exposure **(Fig. 2D)**. These antigens were primarily GAS virulence factors involved in pathogenesis and immune evasion (e.g., SLO, M1, C5AP, SPEB etc.). The amino acid sequence for each of the 72 antigens was compared across 2275 publicly available GAS genomes, which revealed high gene carriage, based on their presence in >90% of all genomes, and high sequence conservation, based on low sequence entropy and gap occurrences. One notable exception was the M1 protein, due to the high sequence variability of the hypervariable region (HVR) **(Fig. 2E)**(25). In summary, similar to pooled samples, the individual antigenomes converged around a small subset of genomically conserved antigens. These profiles were similar across healthy individuals, but were considerably different in patients with GAS bacteremia.

**Fig. 2.**
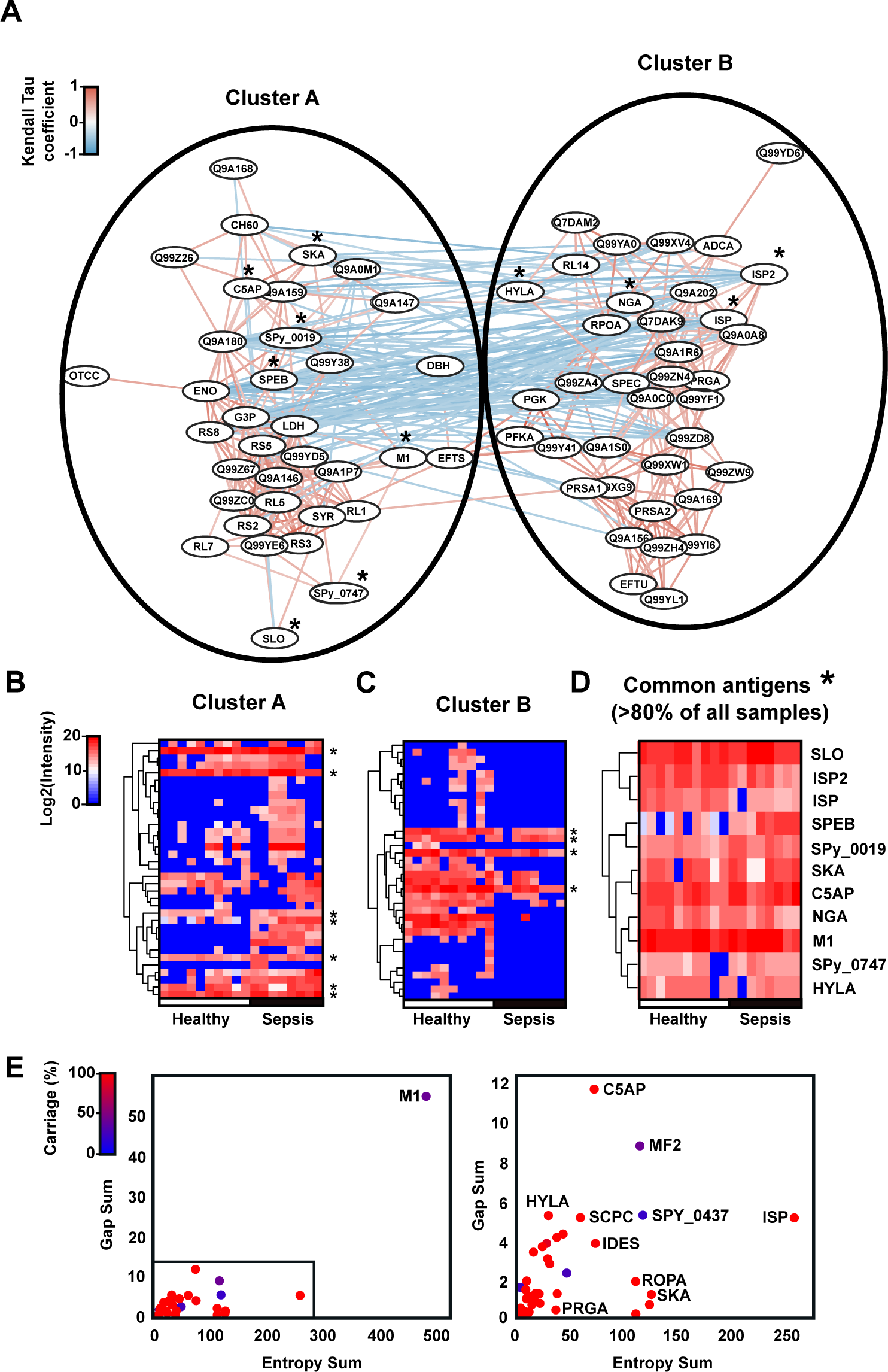
The GAS-specific IgG antigenome across healthy and sepsis individuals. **(A)** Rank correlation network of the 72 GAS antigens identified across different individuals. Nodes represent each identified GAS antigen and the distance between the nodes is defined by edges encoding Kendall tau coefficients for each pairwise comparison. The color of the edges reflects positive (red) or negative (blue) correlation coefficients. To be considered part of the antigenome, the proteins were required to be present in at least two out of three biological replicates, and identified by at least two quantifiable peptides, having at least a two-fold enrichment over the negative control (Xolair, a commercial anti-IgE monoclonal). Pearson correlation clustering of the log2 intensity of the antigens in **(B)** Cluster A, **(C)** Cluster B and **(D)** the 11 common antigens across healthy and sepsis individuals. **(E)** Sequence conservation plots of the 72 antigens based on the analysis of gap frequency, entropy and gene carriage of each protein across 2275 GAS genomes (left), and zoom in plot of the antigens excluding M1 (right). Residues with high conservation have low entropy, whereas residues with low conservation have high entropy. Gaps indicate insertion and deletion in sequences.

### Mapping antigenic sites frequently targeted by circulating GAS-antibodies

The antigenome analysis pointed to a defined set of antigens commonly targeted by GAS antibodies, but polyclonal IgG may bind to one or multiple epitopes within a given antigen, which might lead to different biological outcomes. To determine the epitope landscape of the GAS-specific IgG, we implemented an epitope extraction (EpXT) workflow (**Fig. 3A**). We selected three antigens: M1 and C5AP, identified in Cluster A, and PRGA, identified in Cluster B (**Fig. 2A**). All three proteins were recombinantly expressed and subjected to limited proteolysis to generate partially digested protein regions of different sizes. The partial digests were captured by immobilized IVIG antibodies to isolate antigenic protein regions, which were eluted and quantified by LC-MS/MS. The method was first applied to C5AP, a streptococcal serine peptidase with a multidomain structure: a protease-associated domain (PA-domain), a catalytic domain (Cat-domain), and three consecutively arranged fibronectin-type domains (FN-domains) (**Fig. 3B**)(26). The EpXT analysis identified 17 immunogenic peptides (**Supplemental table 4**). Roughly 70% of the total peptide intensity was associated with the Cat-domain, ∼15% with the FN1-domain, and only ∼5% with the FN2 domain (**Fig. 3B**). Importantly, the interaction of C5AP with IVIG was validated by hydrogen-deuterium exchange mass spectrometry (HDX-MS), which identified two peptide stretches (97-138 aa and 415-466 aa) displaying a significant reduction in deuterium uptake upon incubation with IVIG (**Fig. 3C & Supplemental table 5**). These binding sites partially overlapped with the ones identified by the EpXT-workflow, demonstrating good agreement between the methods, and singling out the Cat-domain as an immunodominant region (**Fig. 3D).**

**Fig. 3:**
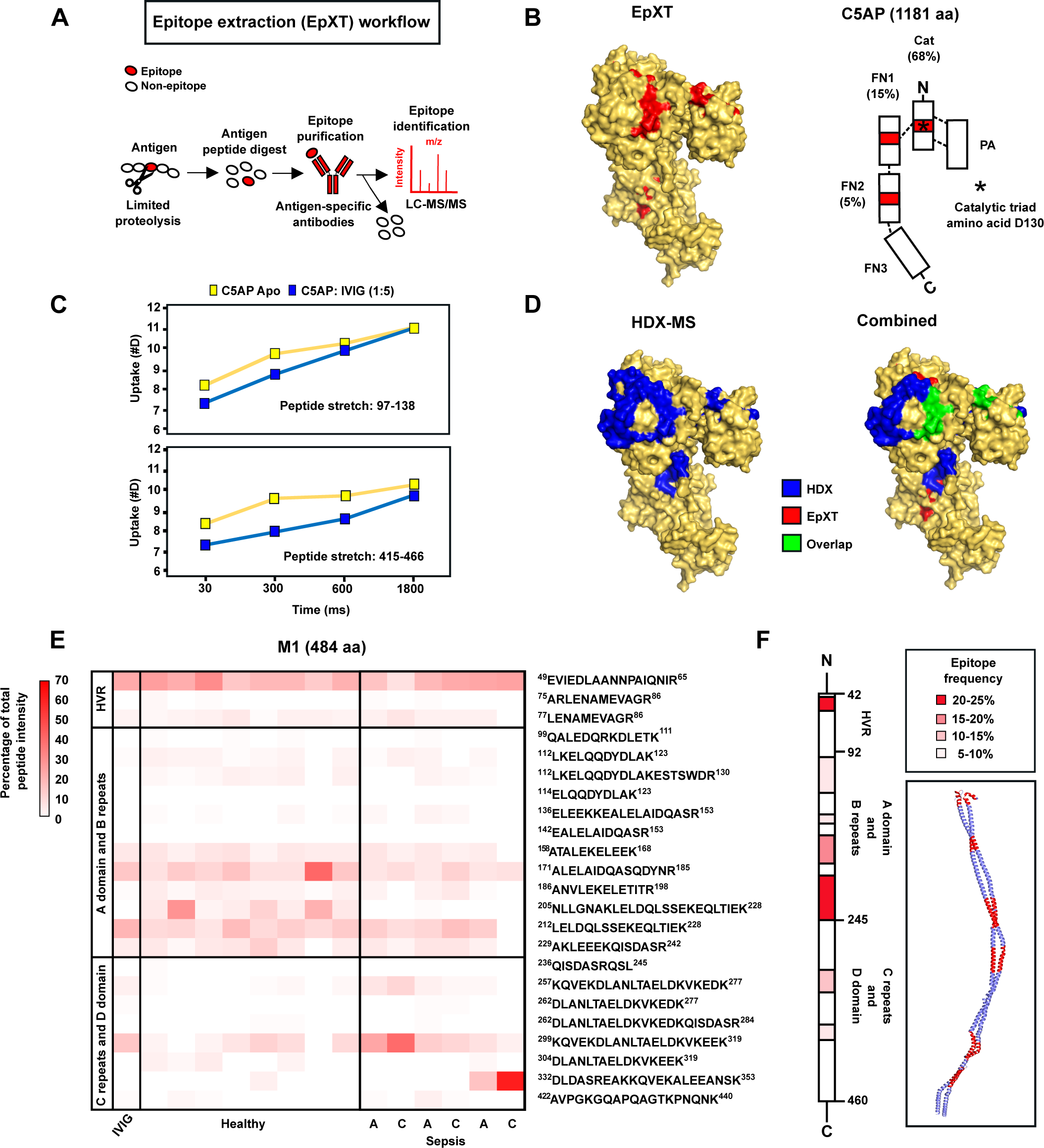
Epitope mapping of GAS antigens. **(A)** Schematic representation of the epitope extraction workflow (EpXT) to identify epitopes on recombinantly expressed antigens. **(B)** Identified peptides (marked red) by EpXT mapped onto the crystal structure of C5AP (left) and their relative intensity (%) is shown on the C5AP cartoon (right). **(C)** Deuterium uptake plots for two peptide stretches of C5AP alone (yellow) and when incubated with IVIG (blue). **(D)** Identified peptides (marked blue) by HDX-MS (left) and overlapping epitopes identified by both HDX-MS and EpXT mapped onto the crystal structure of C5AP (right). **(E)** Heatmap of M1 peptide intensities across IVIG, healthy and sepsis individuals. **(F)** Consensus epitope landscape across all individuals with more than 5% epitope frequency are displayed in the M1 cartoon and the M1 model.

The interaction between IVIG and the M1-protein was also studied by EpXT. M1 is a dimeric coiled-coiled fibrillar protein with an N-terminal HVR, a variable region encompassing the A domain and B repeats, and a constant region comprising the C repeats and D domain. We identified 23 antigenic peptides across the three regions. Roughly 26% of the intensity was associated with the HVR, 52% with the variable region and 22% with the constant region (**Fig. 3E)**. To address whether similar epitopes are recognized by IgG from different individuals, we interrogated plasma from healthy individuals, as well as the paired acute and convalescent plasma from the patients with GAS bacteremia. The individual epitope patterns were consistent with the pattern observed when using IVIG, both in terms of the peptide identities and their relative intensity distributions (**Fig. 3E**). Overall, the epitope profiles were similar across healthy individuals, and between the acute and convalescent plasma of each patient, although peptides from the constant region of M1 tended to be more enriched when using plasma from patients with bacteremia. Averaging the peptide signal across all samples showed that the HVR and the variable region are the most commonly targeted and immunodominant sites **(Fig. 3F)**.

Finally, PRGA, an 873 aa long GAS protein of unknown function was also analyzed by EpXT. Since there is no structure available, a molecular model of PRGA was generated, which predicted an extended coiled-coiled structure with an internal globular domain **(Fig. S4**). A total of 6 peptides were identified, with ∼90% of the intensity associated with the globular domain, thereby also indicating antibody binding to spatially confined and most likely immunodominant regions of PRGA (**Supplemental table 4**). In conclusion, we show that EpXT can map immunodominant antigenic sites on various GAS antigens. For the M1-protein, these sites are conserved across individuals, suggesting common mechanisms of epitope recognition.

### The IgG subclass impacts the ability of anti-M1 antibodies to trigger immune signaling

In addition to neutralization through antigen and epitope recognition, antibodies can elicit protective effector functions that are dependent on other IgG properties, such as the Fc-glycosylation and the IgG subclass distribution(2). To test how GAS antibodies trigger Fc-dependent immune signaling, recombinant M1, C5AP and PRGA were incubated with IVIG and probed for the activation of FcγR-receptor IIa (CD32), a surrogate for ADCP, and IIIa (CD16), a surrogate for ADCC, using luciferase reporter cell assays. Antibodies against all three antigens elicited both CD32 and CD16 activation, with significant variation observed across the antigens (C5AP>PRGA>M1) (**Fig. 4A & 4B**). These differences could not be explained by titers (M1>C5AP>PRGA) (**Fig. 4C**) or glycosylation, as glycoproteomic analysis of the antigen-specific IgG ruled out major differences in the Fc glycan profiles (**Fig. S5A and method section**). However, the LC-MS/MS quantification of the affinity-purified IgG subclasses using proteotypic Fc peptides showed that more IgG of each subclass was pulled down using M1 as a bait, compared to C5AP and PRGA (**Fig. 4D**). The subclass distribution was also skewed and M1-antibodies were more enriched in the IgG2 and IgG3 subclasses, compared to antibodies recognizing the two other antigens.

**Fig. 4:**
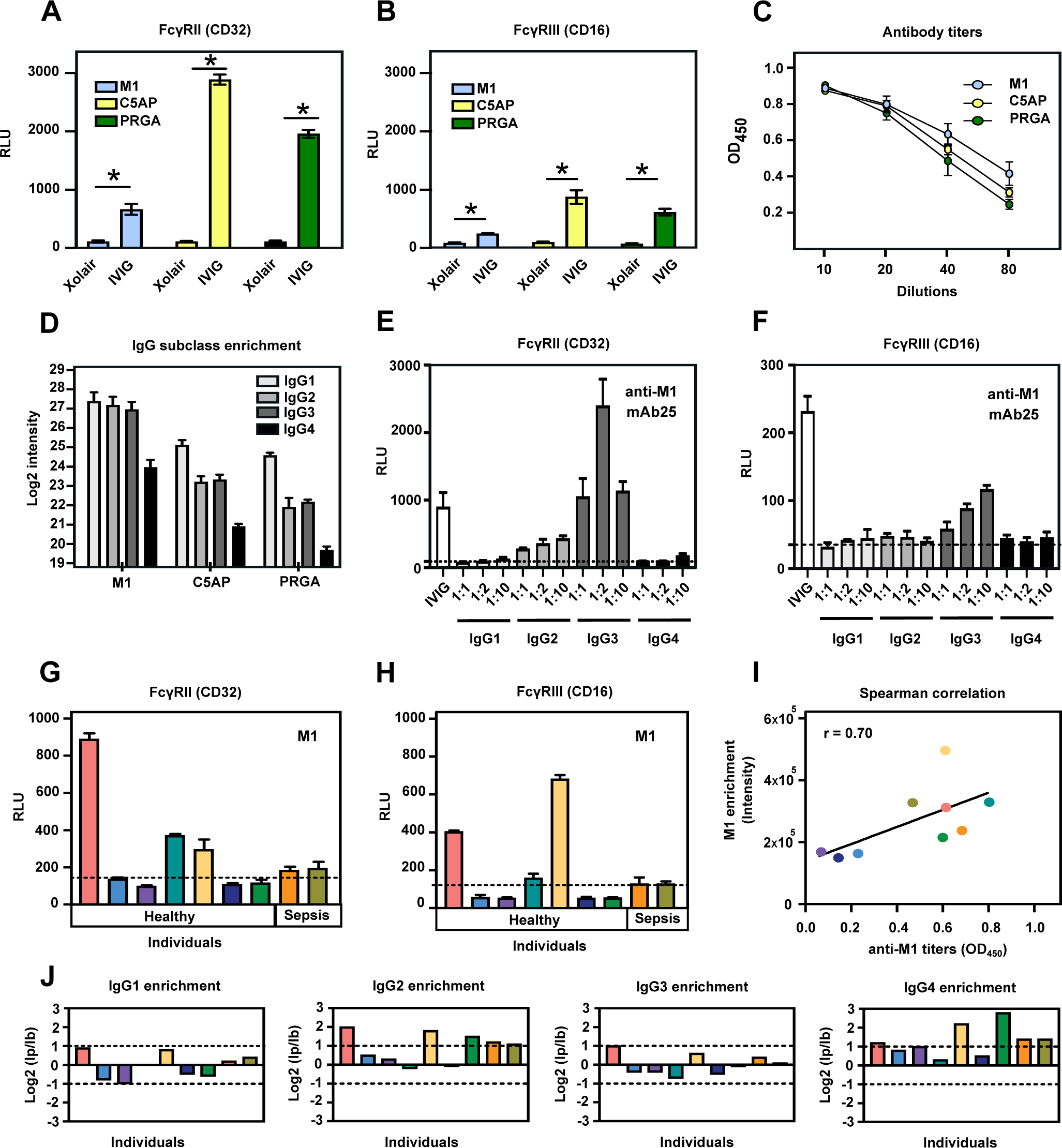
GAS-specific antibodies trigger immune signaling in an antigen- and receptor-specific manner. **(A)** FcγRII (CD32) and **(B)** FcγRIII (CD16) activity assay of M1, C5AP and PRGA specific antibodies present in IVIG. **(C)** Antibody titers in IVIG against M1, C5AP and PRGA. **(D)** Subclass enrichment profiles of antigen-specific IgG in IVIG. **(E)** FcγRII (CD32) and **(F)** FcγRIII (CD16) activity assay of mAB25 in IgG1-4 scaffolds. **(G)** FcγRII (CD32) and **(H)** FcγRIII (CD16) activity assay of M1-specific antibodies across individual plasma samples. **(I)** Correlation plot of M1 enrichment and titers across healthy and infected individuals. **(J)** Subclass enrichment profiles of M1-specific IgG across individuals after normalizing the intensity of pulldowns (I_p_) against the bulk (I_b_). Statistical significance was assessed by two-way ANOVA, * p<0.05. The results are the average of experiments done in triplicates, and reproduced at least three times.

To better determine the impact of the IgG subclass on the ability of anti-M1 antibodies to trigger FcγR-signaling, we took advantage of the monoclonal mAb25 that specifically binds to the M1-protein with high affinity(27). This monoclonal antibody allowed us to rule out potential confounding factors, such as the relative contribution of mixed subclasses and different epitope binding patterns of the M1-specific IgG in IVIG(28). Notably, whereas the mAb25 in IgG1 or IgG4 scaffolds displayed almost no measurable FcγR-receptor activation, swapping to IgG2 or IgG3 both resulted in robust induction of CD32-signaling (**Fig. 4E),** whereas only IgG3 was capable of triggering CD16-signaling (**Fig. 4F**). These results indicate that the proportionally higher levels of IgG2 and IgG3 among the M1 enriched IgG correlate with enhanced FcγR-receptor activation for this antigen.

Next, we investigated whether anti-M1 antibodies isolated from different healthy donors and individuals with GAS bacteremia were also subject to variation regarding their capacity to trigger CD32 and CD16 activation. As shown in **Fig.4G-H**, roughly half of the samples showed Fc-signaling activity over the baseline with substantial variation observed across individuals. In general, responding individuals had higher antibody titers than low responders, which correlated well with their increased ability to pull down M1 from the bacterial fractions **(Fig. 4I)**. However, the IgG subclasses that were enriched varied considerably between individuals **(Fig. 4J)**. Finally, as observed for pooled IVIG samples, glycan analysis revealed no major variation in Fc glycosylation across individuals (**Fig. S5B**). Combined, these results indicate that a direct correlation between the structural properties of the antigen-specific IgG repertoires and FcγR-signaling is challenging to decipher in polyclonal antibody pools. In contrast, using defined monoclonal antibodies demonstrate that the IgG subclass has a major impact on the capacity of anti-M1 antibodies to trigger immune signaling. A final summary of the main structural and functional properties of circulating GAS-specific antibodies uncovered by the systems antigenomics-serology workflow is presented in **Fig. 5**.

**Fig. 5:**
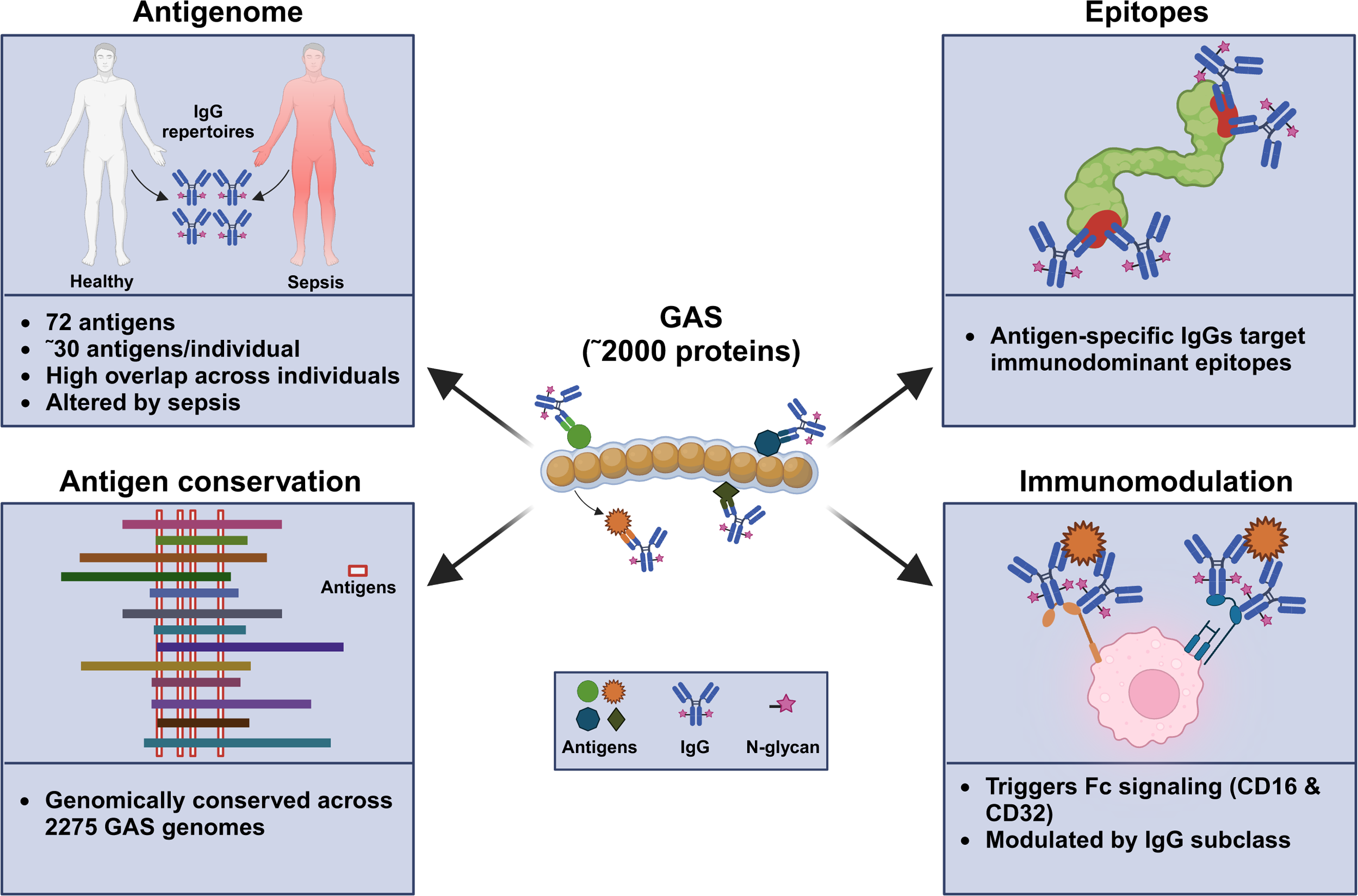
Schematic summary of the GAS-specific IgG antigenome and the structural and functional properties of circulating GAS-specific IgG uncovered by the systems antigenomics-serology pipeline.

## Discussion

In this study we developed a novel MS-centered methodology that couples systems antigenomics to systems serology to dissect the key properties of polyclonal antibody responses directly in human samples, in a reproduceable, high-throughput and flexible manner. The method involves surveying the antigen repertoire targeted by the antibodies using fractionated pools of bacterial proteins and immunologically reactive sera. Once the antigens are identified and ranked, they are recombinantly expressed and the antigen-specific IgG is interrogated as to their structural and functional properties, including epitope repertoires, subclass distribution, Fc glycosylation pattern, and capacity to trigger immune signaling. We applied this methodology to dissect the key features of naturally occurring GAS-specific IgG circulating in adult human plasma.

GAS has been the subject of intensive research due to its high disease burden, broad spectrum of pathogenic mechanisms, and geographically constrained serotype prevalence(25). A roadmap towards a GAS vaccine has recently been outlined by the World Health Organization (WHO) to contain the burden of both local and invasive streptococcal infections, as well as their autoimmune sequelae(29). However, major challenges remain, including the lack of reliable immune correlates of infection and protection, and a poor understanding of the evolution of the immune response against GAS during natural exposures. Notably, the incidence of GAS infections is high in school-age children but typically declines throughout life, which suggests the buildup of protective immunity during the lifetime of an individual(25).

Our data confirm that most adults have circulating IgG against the GAS antigenome, a relatively small subset of streptococcal antigens that are genomically conserved across GAS isolates, and commonly targeted by circulating antibodies across multiple individuals. A typical GAS genome codes for ∼1800 proteins(30, 31) so the finding that the size of the GAS antigenome is on average 30 antigens/individual raises the question why some proteins are more frequently targeted by host antibodies than others. According to our data, the GAS antigenome covers a wide range of molecular structures and functions, ranging from multimeric adhesion proteins, such as the M-protein, to highly specific monomeric proteases, such as C5AP. However, despite its smaller size compared to the expected proteome, the GAS antigenome is enriched in virulence factors. Since GAS is a human-adapted pathogen, it is possible that some of these factors have been targeted by host antibodies as a consequence of an arms race between bacterial virulence and host immunity during evolution. In addition to high genomic conservation, most of these factors are known to directly facilitate the establishment of a successful infection, and may therefore be produced in high amounts during host-pathogen encounters compared to other streptococcal proteins, which in turn might result in greater accessibility to immune and antigen-presenting cells. In addition, the antigenic breath of the GAS-specific antibodies might also be dynamically regulated by the immune status, which would be in line with our finding that the GAS antigenome is different in patients with bacteremia compared to healthy individuals. Although the size of our cohort was rather small to draw firm conclusions, our data suggest that the GAS antigenome is sensitive to ongoing infections, which can be further explored by combining our techniques with larger and more defined clinical cohorts covering different types of streptococcal infections.

Another possibility for the seemingly small size of the GAS antigenome might be due to technical rather than biological reasons. Our antigenomics strategy relies on fractionated protein pools extracted from growing bacteria, and hence changes in the culture conditions might result in altered proteome profiles that would lead to some antigens being missed due to differential expression. Indeed, some virulence factors such as the streptococcal endoglycosidase EndoS was not identified in our screen, despite anti-EndoS antibodies being widely present in human plasma(32). It is therefore possible that both growing conditions (e.g., exponential vs stationary growth phases, presence vs absence of plasma etc.) or even the specific strain used to generate the bacterial protein pools might determine the repertoire of antigens available to the antibodies during the screen. Still, our findings are in line with previous studies using completely different methods, such as protein arrays and surface display technology, which suggests that most of the core antigenome is efficiently captured by our methodology(7, 10, 11, 33). The relatively high agreement between these studies and the antigenome profiling presented in this report, confirms our notion that natural exposure to GAS results in a distinct serological signature dictated by the immunological recognition of a relatively small and well-defined set of streptococcal antigens. As opposed to these previous studies, here we generated libraries of bacterial proteins through biochemical fractionations. This has the advantage of reducing the high costs and expertise associated with surface display and protein array technology, making our strategy more amenable to any biochemical laboratory with access to standard bacterial growth facilities and equipment. Additionally, cellular fractionation allows querying properly folded proteins associated with relevant and immunologically accessible compartments, such as membrane and cell wall proteins, since the actual localization of many proteins might still be difficult to predict using genome mining and reverse vaccinology strategies. Finally, our workflow is flexible and fully automated, and can be exploited to query a wide range of growing conditions and cohorts, as well as being easily adapted to analyze other bacteria.

In addition to analyzing the GAS antigenome, we took a step further and developed approaches to map the epitope landscape of selected GAS antigens. Interestingly, antibody recognition was invariably associated with defined antigen sites or immunodominant regions. Although much is known regarding T-cell immunodominance, the basis for B-cell and antibody immunodominance is less well understood(34). Our detailed dissection of the epitope landscape of the M protein, a promising immunogen for a GAS vaccine, showed that the HVR and the variable region are major interaction sites for naturally occurring GAS antibodies. The variability of the HVR is thought to be the result of selective pressure on the bacteria to escape the immune response, since type-specific antibodies are protective against infections(35, 36). However, previous studies indicated that the HVR might be only weakly immunogenic(37), which contrast with our observation of antibodies binding to the HVR across all samples studied. One possible explanation for this discrepancy is that natural exposure to GAS is often accompanied by a robust induction of the immune response during infection, which might create an appropriate environment for selection of B-cell clones targeting the HVR. These conditions might not be completely phenocopied by immunization studies using laboratory animals.

Finally, our approaches also facilitated the analysis of key attributes of the Fc regions of naturally occurring GAS-specific antibodies. We show that GAS antibodies can engage multiple FcγRs and robustly trigger immune signaling, at least *in vitro*, in an antigen- and Fc receptor-specific manner. The affinity of FcγRs for IgG varies with the Fc structure, in particular the subclass and glycosylation, and enrichment of specific subclasses and pro- or anti-inflammatory glycan structures is often an avenue exploited by the host immunity to modulate the affinity of these interactions during infection and vaccination(2). In the case of M-specific antibodies, signaling varied across individuals, correlated with IgG titers, and was modulated by the IgG subclass distribution. Complement deposition and opsonophagocytosis mediated by type-specific antibodies recognizing the HVR region of the M protein is a well-known correlate of protection against GAS-infections(36). However, whether FcγRs also contribute to protection is less clear. FcγRs are important orchestrators of immunomodulation and protection against many pathogens, aiding in phagocytosis and immune cell degranulation. Whether these mechanisms are also relevant for GAS infections *in vivo* remains to be determined.

## Data availability statement

The mass spectrometry and HDExaminer analysis files have been deposited to the MassIVE repository with the dataset identifier MSV000093310.

## Acknowledgements

J.M. is a Wallenberg academy fellow (KAW 2017.0271) and is also funded by the Swedish Research Council (Vetenskapsrådet, VR) (2019-01646 and 2018-05795), the Wallenberg foundation (WAF grant number 2017.0271), and Alfred Österlunds Foundation. L.M. is funded by the Swedish Research Council (Vetenskapsrådet, VR) (VR-2020-02419) and Alfred Österlunds Foundation. This work was supported by generous funding from NIGMS (R35 GM119850 to N.E.L.) and the Novo Nordisk Foundation (NNF20SA0066621 to N.E.L.).

## Materials and methods

### Patient enrolment and sample collection

The sampling of patients with bacteremia was approved by the regional ethics committee of Lund University, (2016/939, with amendment, 2018/828). Oral and written consents were obtained from included participants. During 2018-2020, four patients with GAS bacteraemia in Region Skåne, Sweden, were enrolled in the study. Acute sera were collected within five days after hospital admission, and convalescent sera were collected after 4-6 weeks. Information on the four included patients is given by *de Neergaard et.al* (24). Citrated blood samples were collected from 10 healthy donors. Platelet-poor plasma was prepared by centrifugation at 2000 x g for 10 min and stored at −80°C until use. Ethical approval was obtained from the local ethics committee (approval 2015/801).

### Biochemical fractionation of GAS proteins

A single colony of the M1 GAS serotype SF370 was precultured in Todd-Hewitt broth supplemented with 0.6% yeast extract (THY) at 37°C and 5% CO_2_ for 16-18 hr (OD_620nm =_ 0.8) and then the bacteria was sub-cultured in either protein reduced THY broth or regular THY broth. The protein reduced THY broth was prepared by passing THY broth through a 0.22-µm-pore-size-filter and then filtered using a 10-kDa molecular mass cut-off. For the secreted fractions, bacteria grown in protein reduced THY broth at 37°C & 5% CO_2_ till the mid-logarithmic phase (OD_620nm_=0.4-0.5) were harvested at 3000 g for 15min at 4°C and the culture supernatant was filtered using a 0.22-µm-pore-size-filter unit. The filtered supernatant was concentrated using ice cold 1X phosphate buffered saline (PBS) using amicon ultracel 10kDa molecular weight cutoff centrifugal filtration unit (Millipore) at 4000g for 15 minutes (min) and stored in −20°C until further use.

For the cell wall and membrane fractions, bacteria were sub-cultured in regular THY broth at 37°C & 5% CO_2_ to mid-logarithmic phase (OD_620nm_ 0.4-0.5) and the cells were harvested at 3000 g for 15min at 4°C. The bacterial pellets were kept on ice for a brief period of 5 min followed by resuspension in 5 ml chilled TES buffer (50mM Tris-HCl, 1mM EDTA, 20% sucrose (w/v) sucrose, pH 8.0) containing 1mM phenylmethylsulfonyl fluoride (PMSF, Roche) at 7560g for 20 minutes at 4°C. For bacterial cell wall lysis, 1.15ml of ice-cooled mutanolysin mix (1ml TES buffer, 100 µl lysozyme (100mg/ml in TES), 50 µl mutanolysin (Sigma-Aldrich, 5000U/ml in 0.1M K_2_HPO4, pH 6.2) was added to the cells for 2 hr at 37°C at 200rpm shaking. Cells were then centrifuged at 14000g for 5min and the resulting supernatant had the cell wall fractions which was stored in −20°C until further use.

To isolate membrane proteins, the cell pellets were washed twice in 1 ml HEPES-buffer at 3500g for 5min and the cells were then dissolved in 1% HEPES. 2 µl of 0.5 µG/µL trypsin was added to cells for 60 min at 37°C at 500 rpm to shave off the membrane proteins and the reaction was stopped by incubating the cells on ice for 2 min before centrifuging them at 1000g for 15min at 4°C. The supernatant containing the membrane proteins was then collected and stored in −20°C. Cell pellets were further treated with RIPA lysis buffer for 15 minutes and centrifuged 3500g for 5min to collect the intracellular fractions.

### IgG immunoblotting

Secreted, cell wall, and membrane GAS protein fractions were separated on SDS-PAGE (Criterion TGX Gels, 4%–20% precast gels, Bio-Rad) and proteins were transferred to PVDF membranes using the trans-blot turbo transfer system (BioRad) according to the manufacturer’s instructions. The membranes were blocked with 5% bovine serum albumin (BSA) in PBST (PBS + 0.05 % Tween 20) for 1 hr at 37°C, followed by incubation with IVIG (Octagam) (1:100) and pooled human plasma (1:10, Innovative research) overnight at 4°C. After washing, the membranes were incubated with protein G-HRP conjugate (1:3000, Bio-Rad) for 1 hr at 37°C. The membranes were then developed using clarity western ECL substrate (Bio-Rad) and visualized in the ChemiDoc MP Imaging System (Bio-Rad).

### Enzyme-linked immunosorbent (ELISA) assay

To measure GAS-specific antibodies, 96 well Nunc microtiter plates were coated with 100 µl of recombinant M1, C5AP and PRGA (5µg/ml) overnight at 4°C followed by PBST (PBS + 0.05 % Tween 20) wash. Plates were blocked with 2% BSA (100 µl/well) in PBST for 30 min at 37°C. After washing with PBST, IVIG (1:100) and plasma (1:10) was added in dilution series in triplicates and incubated at 37°C for 1 hr and then washed with PBST. 100 µl/ well of protein G-HRP conjugate (1:3000, Bio-Rad) in PBS was added and incubated for 1 hr at 37°C and then washed with PBST. The reaction was developed using 100 µl/well ABTS (20 ml Sodium Citrate pH 4.5 + 1ml ABTS + 0.4 ml H_2_O_2_) for 30 min and the OD was measured at 450 nm.

### FcγR-luciferase reporter cell assay

Jurkat-Lucia NFAT-CD16 (FcγRIII) and CD32 (FcγRII) cells (InvivoGen) were used to probe the ability of antigen specific IgG to trigger antibody-dependent cellular cytotoxicity (ADCC) and antibody-dependent cell-mediated phagocytosis (ADCP). Nunclon delta surface plates (Thermo Scientific) were coated with 100 µl of 5 µg/ml of M1, C5AP and PRGA overnight at 4°C followed by 1XPBS wash. 100 µl of different antibody sources (100 µg/100 µl) *i.e.,* IVIG (1µl of IVIG diluted with PBS to a final volume of 100 µl), Xolair (10 µl of Xolair diluted with PBS to a final volume of 100 µl) and human plasma (10µl of human plasma diluted with PBS to a final volume of 100 µl) were added and incubated for 1hr at 37°C. After 1XPBS wash, 200 µl of CD16 and CD32 cells (100,000 cells/100 µl) in IMDM with 10% heat-inactivated fetal bovine serum (FBS) and Pen-Strep (100 U/ml-100 µg/ml) were respectively added and incubated at 37°C for 6hr. After a brief centrifugation for 10 min at 150g, 20 µl of the supernatant was added to 50 µl of QUANTI-Luc (InvivoGen) in opaque microtiter plates and the luciferase activity was measured in luminometer.

### Affinity purification of bacterial antigens

IgGs from different sources were purified in a 96 well plate (Greiner) using the Protein G AssayMAP Bravo (Agilent) system, according to the manufacturer’s instructions. Briefly, 1µl of IVIG (Octagam), 10 µl of Xolair (Omalizumab) and 10µl of human plasma was diluted with PBS to a final volume of 100 µl and then applied to pre-equilibrated Protein G columns. Columns were washed with PBS, before applying a pool of 100µg secreted, 100µg cell wall and 100µg membrane fractions. The antigen-antibody complex was then eluted in 0.1M glycine (pH=2) and the final pH was neutralized with 1M Tris, and saved until further use. The proteins were denatured using 8 M urea solution and 5 mM Tris(2-carboxyethyl) phosphine hydrochloride (TCEP) was then added for 60 min at 37°C to reduce the disulfide bonds followed by alkylation with 10 mM iodoacetamide in the dark at room temperature for 30 min. 100 mM ammonium bicarbonate was added followed by the addition of 0.5 µG/µL sequencing-grade trypsin (Promega) for protein digestion at 37°C for 18 h. The activity of trypsin was inhibited by dropping the pH to 2-3 by the addition of 10% trifluoroacetic acid (TFA, Sigma). The samples were loaded on Evosep tips to separate the digested peptides using nanoflow reversed-phase chromatography with an Evosep One liquid chromatography (LC) system (Evosep One) and analyzed on timsTOF Pro mass spectrometer (Bruker Daltonics).

### Antigen-specific IgG pulldowns

Antigen specific IgG was purified from IVIG and human plasma in a 96 well plate setup according to the manufacturer’s instructions. IgG from 100 µl of human plasma (∼100 µg/100 µl) was pre-enriched using the Protein G AssayMAP Bravo (Agilent) technology as described above. Eluted IgG was diluted to a final volume of 500 µl with 1XPBS and then buffer exchanged in 50K centrifugal filters (Amicon Ultra-0.5 ml, Merck) for 10 min at 14000g and was finally resuspended in 100 µl of 1XPBS and treated as bulk IgG enriched from human plasma. For the antigen specific pulldowns, 20 µg of recombinantly expressed M1, C5AP and PRGA with streptavidin tag were immobilized on pre-equilibrated AssayMAP Streptavidin columns (Agilent Technologies). Columns were washed with 1XPBS and then 100 µl of IVIG (1µl of IVIG diluted with 1XPBS to a final volume of 100 µl) and 90 µl of pre-enriched IgG from human plasma was applied followed by 1XPBS wash. Elution was done using 100 µl of 0.1M glycine (pH=2) and the final pH was neutralized with 20 µl of 1M Tris. 120 µl of antigen specific IgG, 1 µl of IVIG and 10 µl of bulk IgG from human plasma was diluted to a final volume of 220 µl using 100 mM ammonium bicarbonate, followed by digestion to peptides using 1 µg trypsin at 37°C for 18 hr and the digestion was stopped using 20% TFA (Sigma) to pH 2 to 3. Peptide clean-up was performed using AssayMAP C18 columns (Agilent Technologies) according to manufacturer’s protocol. Samples were dried using vacuum concentrator (Eppendorf) and resuspended in 20 µl .1% formic acid (FA, Fisher Chemical) followed by a brief sonication for 5 min before analyzing on a Q Exactive HF-X mass spectrometer (Thermo Scientific).

### Epitope extraction (EpXT)

To benchmark the EpXT workflow Pierce Protein G magnetic beads (Thermo Scientific) were used. For IgG enrichment, 50µl of protein G beads was washed with 1XPBS, before 1µl of IVIG (Octagam) diluted with PBS to a final volume of 100 µl (100µg/100µl) was added and incubated for 1 hr followed by 1XPBS wash. 10µg of recombinant C5AP and PRGA was trypsinized with 1µg of trypsin (Sequencing Grade Modified Trypsin, Promega) for 15 min at 37°C and the trypsin activity was inhibited by incubating at 100°C for 5 min. The peptide digest was then incubated with protein G enriched IgG for 1 hr and then washed with 1XPBS before eluting with 100 µl of 0.1 M glycine (pH=2) and the pH was finally neutralized with 1M Tris. Peptide clean-up was performed on Evosep columns as mentioned above before analyzing on a timsTOF Pro mass spectrometer (Bruker).

For M1 EpXT analysis IgGs from IVIG and human plasma were purified in a 96 well plate (Greiner) using the Protein G AssayMAP Bravo (Agilent) system. 1µl of IVIG (Octagam) and 10µl of human plasma was diluted with PBS to a final volume of 100 µl (100µg/100µl) and then applied to pre-equilibrated Protein G columns. Columns were washed with PBS, before applying the M1 peptide digest. The M1 peptide digest was prepared by incubating 10 µg of M1 with .1 µg trypsin at 37°C for 15 min followed by a brief incubation at 100°C for 5 min. After PBS wash, the M1 peptide-antibody complex was then eluted in 0.1M glycine (pH=2) and the final pH was neutralized with 1M Tris. Peptide clean-up was performed on Evosep columns according to the manufacturer instructions before analyzing on a timsTOF Pro mass spectrometer (Bruker Daltonics).

### LC-MS/MS proteome analysis

Peptide analysis using data-dependent mass spectrometry (DDA-MS) was performed on a Q Exactive HFX instrument (Thermo Scientific) connected to an Easy-nLC 1200 system (Thermo Scientific). An Easy-Spray column (50-cm, column temperature of 45°C, Thermo Scientific) operated at a maximum pressure of 8 × 10^7^ Pa separated the peptides, and a linear gradient of 4% to 45% acetonitrile in aqueous 0.1% formic acid was run for 65 min. One full MS scan (resolution of 60,000 for a mass range of 390 to 1210, automatic gain control = 3e6) was followed by MS/MS scans (resolution of 15,000, automatic gain control = 1e5) of the 15 most abundant signals. 2 m/z isolation width was set for precursor ions and higher-energy collisional-induced dissociation (HCD) at a normalized collision energy of 30 was used for fragmentation. For peptide analysis on timsTOF Pro, a 30 SPD method (gradient length = 44 min) was used for the separation using an 8 cm x 150 μm Evosep column packed with 1.5 μm ReproSil-Pur C18-AQ particles. A captive source coupled to Evosep One was mounted on the timsTOF Pro mass spectrometer (Bruker Daltonics) which was operated in DDA PASEF mode with 10 PASEF scans per acquisition cycle with accumulation and ramp times of 100 ms each. The target value was set to 20,000, dynamic exclusion was set to 0.4 min and singly charged precursors were excluded. The isolation width was 2 Th for m/z < 700 and 3 Th for m/z>800.

### Glycoproteomics analysis

Purified IgG glycopeptides were analyzed on a Q Exactive HF-X mass spectrometer (Thermo Fisher Scientific) connected to an EASY-nLC 1200 ultra-HPLC system (Thermo Fisher Scientific). Peptides were trapped on precolumn (PepMap100 C18 3 μm; 75 μm × 2 cm; Thermo Fisher Scientific) and separated on an EASY-Spray column (Thermo Fisher Scientific). Mobile phases of solvent A (0.1% formic acid), and solvent B (0.1% formic acid, 80% acetonitrile) were used to run a linear gradient from 4 to 45% over 60 min. MS scans were acquired in data-dependent mode with the following settings, 60,000 resolution @ m/z 400, scan range m/z 600-1800, maximum injection time of 200 ms, stepped normalized collision energy (SNCE) of 15 and 35%, isolation window of 3.0 m/z, data-dependent HCD-MS/MS was performed for the ten most intense precursor ions.

### Hydrogen-deuterium exchange mass spectrometry (HDX-MS)

The HDX-MS analysis was made using automated sample preparation on a LEAP H/D-X PAL™ platform interfaced to an LC-MS system, comprising an Ultimate 3000 micro-LC coupled to an Orbitrap Q Exactive Plus MS. HDX was performed on 1.2 mg/ml C5AP and IVIG (8 mg/mL), in 1X PBS, at a ratio of 1:2 and 1:5 in one continuous run, with runs of the apo state made in between the interaction runs, in total 4 replicate runs were made for the apo state, for the pAb interaction states triplicate samples were run. 5 µl HDX samples were diluted with 25 µl 20 mM PBS, pH 7,4 or HDX labelling buffer of the same composition prepared in D_2_O, pH_(read)_ 7.0. The HDX labelling was carried out for t = 0, 30, 300, 600 and 1800s at 4°C. The labelling reaction was quenched by dilution of 30 µl labelled sample with 30 µl of 1% TFA (Sigma), 0.4 M TCEP (Sigma), 4 M Urea (Sigma), pH 2.5 at 1°C. 60 µl of the quenched sample was directly injected and subjected to online pepsin digestion at 4°C (in-house immobilized pepsin column, 2.1 x 30 mm). The online digestion and trapping were performed for 4 minutes using a flow of 50 µL/min 0.1 % FA (Sigma), pH 2.5. The peptides generated by pepsin digestion were subjected to on-line SPE on a PepMap300 C18 trap column (1 mm x 15mm) and washed with 0.1% FA (Sigma) for 60s. Thereafter, the trap column was switched in-line with a reversed-phase analytical column, Hypersil GOLD, particle size 1.9 µm, 1 x 50 mm, and separation was performed at 1°C using a gradient of 5-50 % B over 8 minutes and then from 50 to 90% B for 5 minutes, the mobile phases were 0.1 % FA (A) and 95 % acetonitrile/0.1 % FA (B). Following the separation, the trap and column were equilibrated at 5% organic content, until the next injection. The needle port and sample loop were cleaned three times after each injection with mobile phase 5% methanol (MeOH) / 0.1% FA, followed by 90% MeOH / 0.1% FA and a final wash of 5% MeOH / 0.1% FA. After each sample and blank injection, the Pepsin column was washed by injecting 90 µL of pepsin wash solution 1% FA / 4 M urea / 5% MeOH. In order to minimize carry-over a full blank was run between each sample injection. Separated peptides were analysed on a Q Exactive Plus MS, equipped with a HESI source operated at a capillary temperature of 250 °C with sheath gas 12, Aux gas 2 and sweep gas 1 (au). For HDX analysis MS full scan spectra were acquired at 70K resolution, automatic gain control = 3e6, Max IT 200ms and scan range 300-2000. For identification of generated peptides separate undeuterated samples were analysed using data dependent MS/MS with HCD fragmentation.

### Proteomics data analysis

The DDA data was analysed in MaxQuant (version 2.0.3.0). The protein database used for the searches were *Homo sapiens proteome* (UniProt proteome identifier UP000005640), GAS proteome (UniProt proteome identifier UP000000750) compiled with common contaminants from other species in-house. Carbamidomethyl (C) modification was set as fixed modification and oxidation (M) and acetyl (Protein N-term) was set to variable modification. 1% protein false discovery rate (FDR) was allowed and match between runs was enabled. The LFQ intensities reported by MaxQuant was used for analysis. The resulting DDA data sets were analyzed in Perseus (version1.6.15.0 and 2.0.7.0) and R studio (version-4.2.0). Both side t-test with a FDR of 0.05 was used for volcano plot analysis.

### Antigenome network analysis

All statistical methods were implemented using Python (version 3.6.10). Antigen intensities across IVIG, pooled human plasma, healthy donor plasma, and acute and convalescent phase plasma were scaled and ranked. First the antigen-by-antigen Kendall Tau measure was made for correspondence in antigen presentation. Second the sample-by-sample Kendall Tau measure was made for correspondence in sample antigen profiles. The Benjamini-Hochberg procedure was used to control for a FDR of <0.10. The significant Kendall Tau measures formed a network where nodes are defined by antigens and edges between nodes are the Kendall Tau measure. Visualization and analysis of the network layers were conducted through Cytoscape.

### HDX-MS data analysis

PEAKS Studio X Bioinformatics Solutions Inc. (BSI, Waterloo, Canada) was used for peptide identification after pepsin digestion of undeuterated samples. The search was done on a FASTA file with only the RDB sequence, search criteria was a mass error tolerance of 15 ppm and a fragment mass error tolerance of 0.05 Da, allowing for fully unspecific cleavage by pepsin. Peptides identified by PEAKS with a peptide score value of log P > 25 and no modifications were used to generate peptide lists containing peptide sequence, charge state and retention time for the HDX analysis. HDX data analysis and visualization was performed using HDExaminer, version 3.1.1 (Sierra Analytics Inc., Modesto, US). The analysis was made on the best charge state for each peptide, allowed only for EX2 and the two first residues of a peptide was assumed unable to hold deuteration. Due to the comparative nature of the measurements, the deuterium incorporation levels for the peptic peptides were derived from the observed relative mass difference between the deuterated and non-deuterated peptides without back-exchange correction using a fully deuterated sample (38). As a full deuteration experiment was not made full deuteration was set to 75% of maximum theoretical uptake. The presented deuteration data is the average of all high and medium confidence results. The allowed retention time window was ± 0.5 minute. The spectra for all time points were manually inspected; low scoring peptides, obvious outliers and any peptides where retention time correction could not be made consistent were removed.

### Gene carriage, entropy and gap analysis for GAS antigenome

A Basic Local Alignment Search Tool database (BLAST version 2.12) was built on all *Streptococcus pyogenes* genomes available in The Bacterial and Viral Bioinformatics Resource Center (BV-BRC, as of 2023-03-28). The sequences of all the antigens were separately searched towards the database using BLASTp, and all hits covering more than 70% of the query sequence were extracted and multiple sequence alignment (MSA) was generated with MUSCLE (version 3.8.1551). The Shannon entropy and frequency of gaps were calculated, based on the MSA to indicate the level of conservation within a group of sequences.

