## Supplemental Figures for "Dissecting the properties of circulating IgG against Group A Streptococcus through a combined systems antigenomics-serology workflow"

* Equal contribution

### Supplementary figure legends

**Fig. S1: Cellular localization and network analysis of the GAS proteome. (A)** Schematic summary showing the bacterial growth conditions and different steps involved in the fractionation method to obtain secreted, cell wall and membrane fractions. **(B)** Bar plots for GAS proteins (SPEB, SLO, ENO, PRGA, M1 and C5AP) representing the difference in their abundances based on LFQ intensity across secreted (S), cell wall (CW) and membrane fractions. STRING functional analysis of protein networks of proteins identified in **(C)** membrane, **(D)** cell wall and **(E)** secreted fraction.

**Fig. S2: Reproducibility of the antigen identification workflow.** Distribution of antigens across three replicates identified by IgG from **(A)** IVIG and **(C)** human plasma. Pearson correlation plots of the intensity of antigens depicting strong correlation between the replicates for **(B)** IVIG and **(D)** human plasma.

**Fig. S3: GAS antigen enrichment and titers across different individuals. (A)** C5AP and **(C)** PRGA enrichment across healthy and sepsis individuals. **(B)** anti-C5AP and (**D)** anti-PRGA IgG titers for healthy and sepsis individuals.

**Fig. S4: Epitope mapping of PRGA. (A)** Identified epitopes (marked red) by EpXT displayed on the PRGA model. **(B)** Relative peptide intensity (%) of the identified epitopes mapped onto a PRGA cartoon.

**Fig. S5: Quantification of the glycosylation patterns of different IgG Fc subclasses across antigens and individuals.** (A) Glycopeptide analysis of main IgG glycoforms modifying antigen-specific antibodies in IVIG directed towards streptococcal M1, C5AP and PRGA. (B) Glycopeptide analysis of main IgG glycoforms modifying antigen-specific antibodies derived from multiple individual plasma and directed towards streptococcal M1. The intensity of each glycoform is presented as a percentage of the total glycopeptide intensity.

### Supplementary table legends

**Supplementary table 1: Proteins identified in secreted, cell wall, membrane and intracellular GAS fractions.**

**Supplementary table 2: Significantly enriched antigens by IgG from IVIG and pooled human plasma.**

**Supplementary table 3: GAS antigens identified in IVIG, pooled human plasma, healthy individuals and sepsis individuals and their corresponding LFQ intensity.**

**Supplementary table 4: Identified epitopes, their length, start position, end position and corresponding intensity for C5AP, PRGA and M1.**

**Supplementary table 5: Experimental and uptake details for all observed peptides in HDX-MS analysis.**

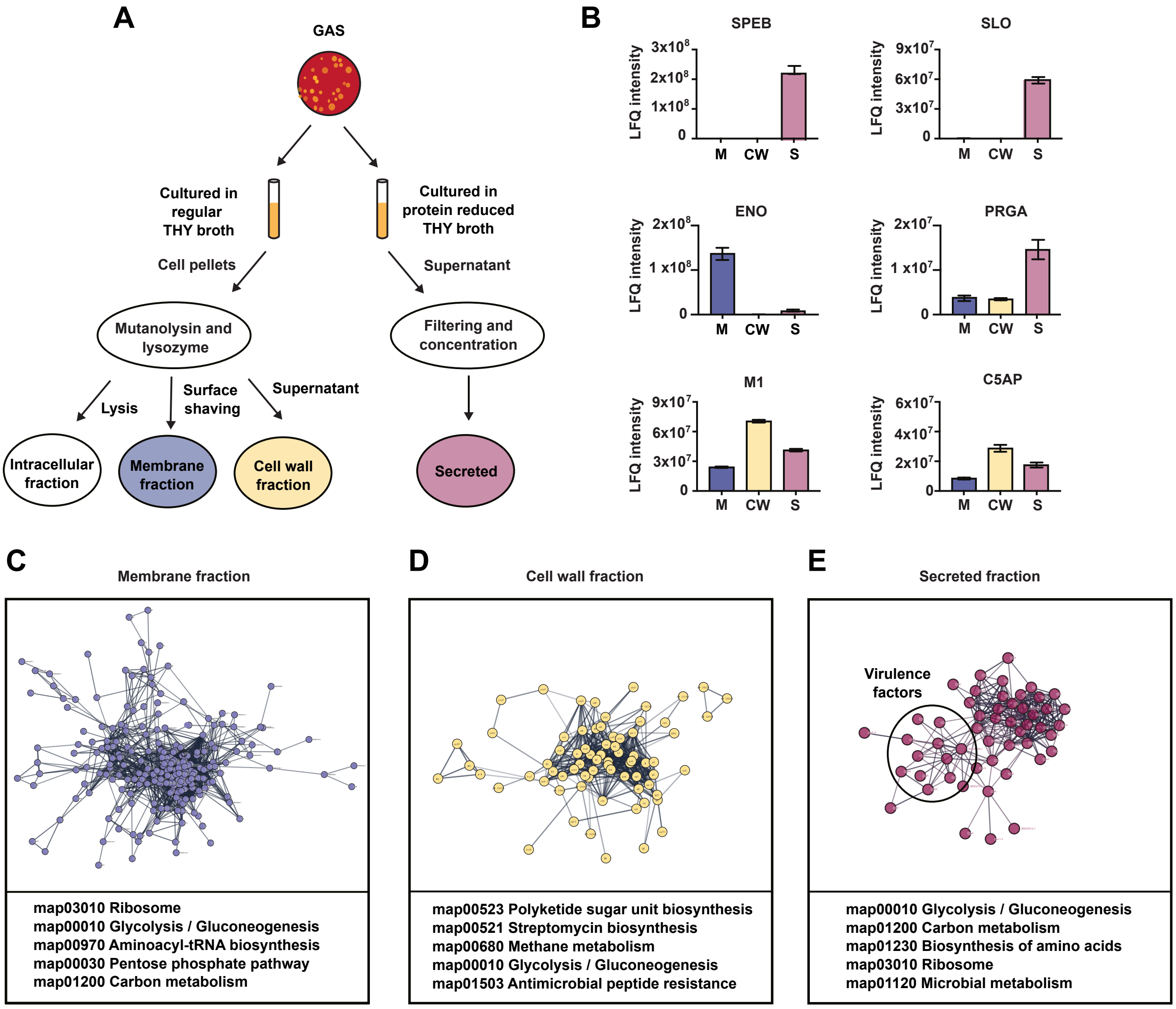

A
GAS
Cultured in regular THY broth
Cultured in protein reduced THY broth
Cell pellets
Supernatant
Mutanolysin and lysozyme
Filtering and concentration
Lysis
Surface shaving
Supernatant
Intracellular fraction
Membrane fraction
Cell wall fraction
Secreted
B
SPEB
SLO
ENO
PRGA
M1
C5AP
LFQ intensity
M
CW
S
C
Membrane fraction
map03010 Ribosome
map00010 Glycolysis / Gluconeogenesis
map00970 Aminoacyl-tRNA biosynthesis
map00030 Pentose phosphate pathway
map01200 Carbon metabolism
D
Cell wall fraction
map00523 Polyketide sugar unit biosynthesis
map00521 Streptomycin biosynthesis
map00680 Methane metabolism
map00010 Glycolysis / Gluconeogenesis
map01503 Antimicrobial peptide resistance
E
Secreted fraction
Virulence factors
map00010 Glycolysis / Gluconeogenesis
map01200 Carbon metabolism
map01230 Biosynthesis of amino acids
map03010 Ribosome
map01120 Microbial metabolism

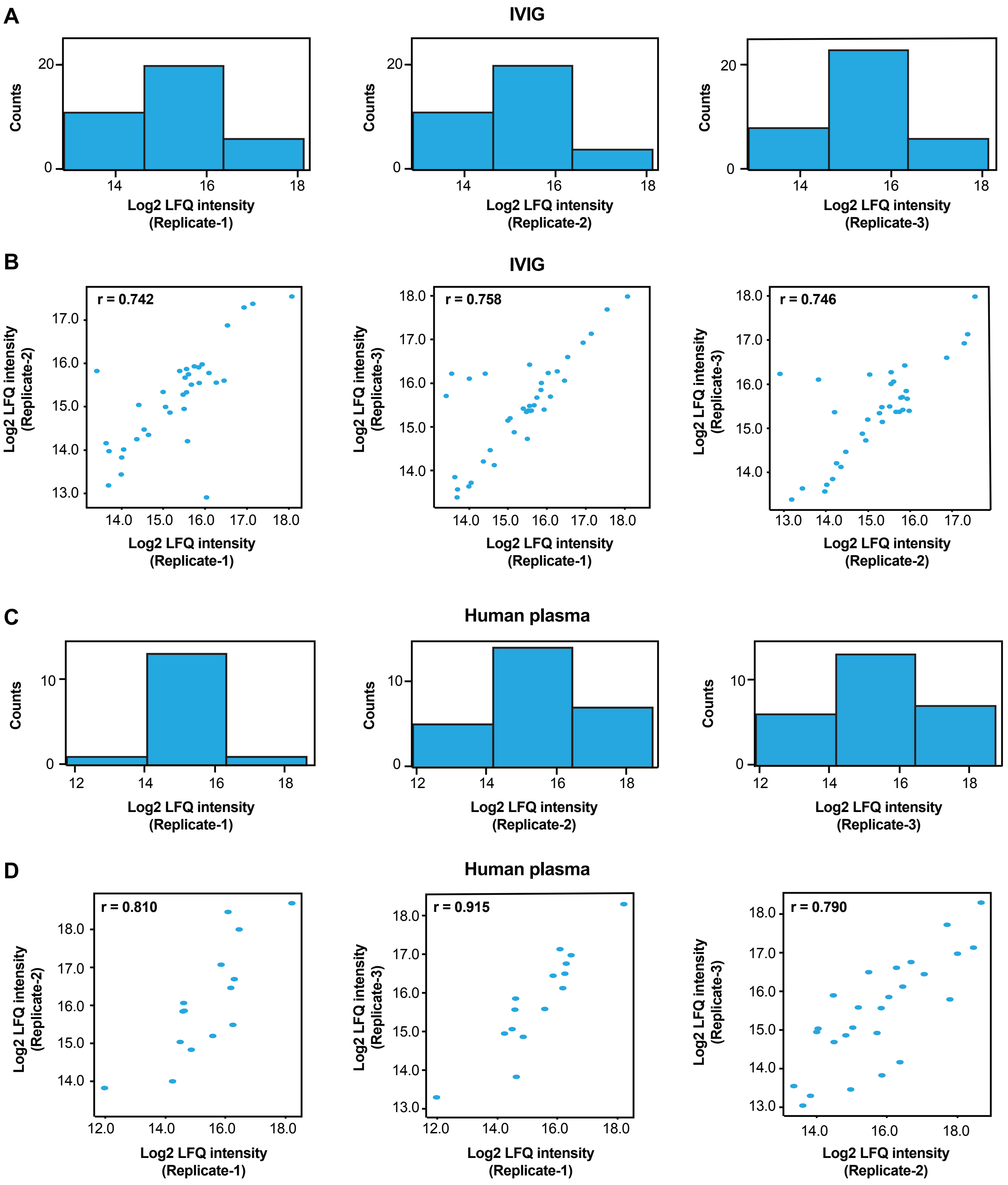
A
IVIG
Counts
Log2 LFQ intensity (Replicate-1)
Log2 LFQ intensity (Replicate-2)
Log2 LFQ intensity (Replicate-3)
B
IVIG
r = 0.742
r = 0.758
r = 0.746
Log2 LFQ intensity (Replicate-1)
Log2 LFQ intensity (Replicate-2)
Log2 LFQ intensity (Replicate-3)
C
Human plasma
Counts
Log2 LFQ intensity (Replicate-1)
Log2 LFQ intensity (Replicate-2)
Log2 LFQ intensity (Replicate-3)
D
Human plasma
r = 0.810
r = 0.915
r = 0.790
Log2 LFQ intensity (Replicate-1)
Log2 LFQ intensity (Replicate-2)
Log2 LFQ intensity (Replicate-3)

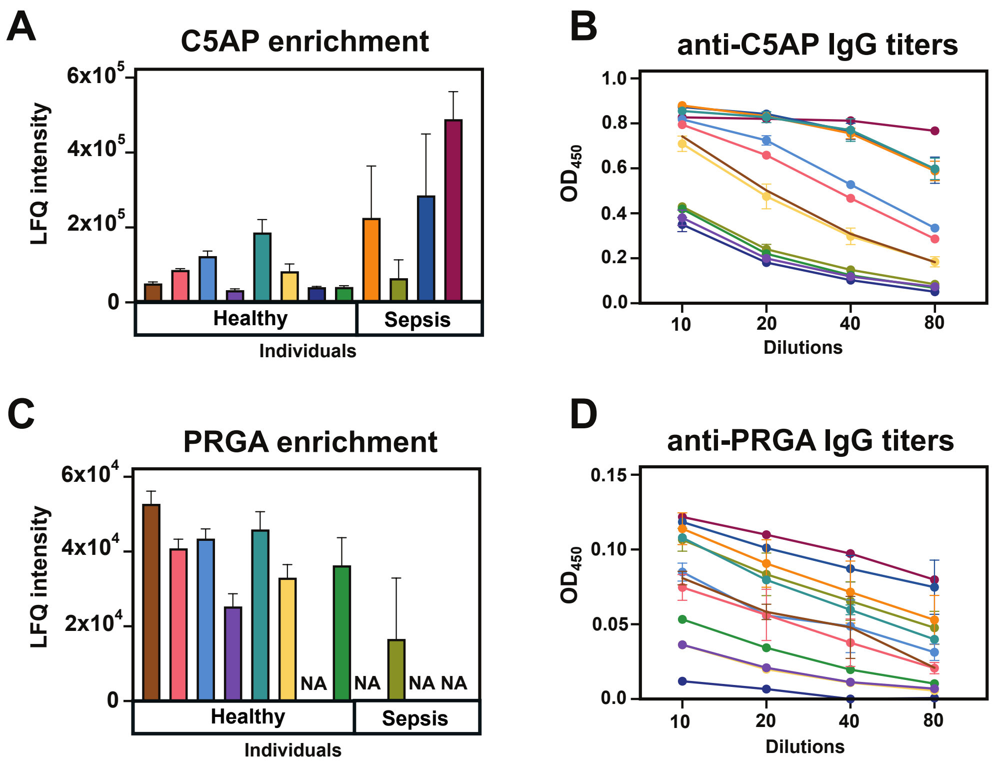
A
C5AP enrichment
LFQ intensity
6x10^5
4x10^5
2x10^5
0
Healthy
Sepsis
Individuals
B
anti-C5AP IgG titers
OD450
1.0
0.8
0.6
0.4
0.2
0.0
10
20
40
80
Dilutions
C
PRGA enrichment
LFQ intensity
6x10^4
4x10^4
2x10^4
0
NA
NA
NA NA
Healthy
Sepsis
Individuals
D
anti-PRGA IgG titers
OD450
0.15
0.10
0.05
0.0
10
20
40
80
Dilutions

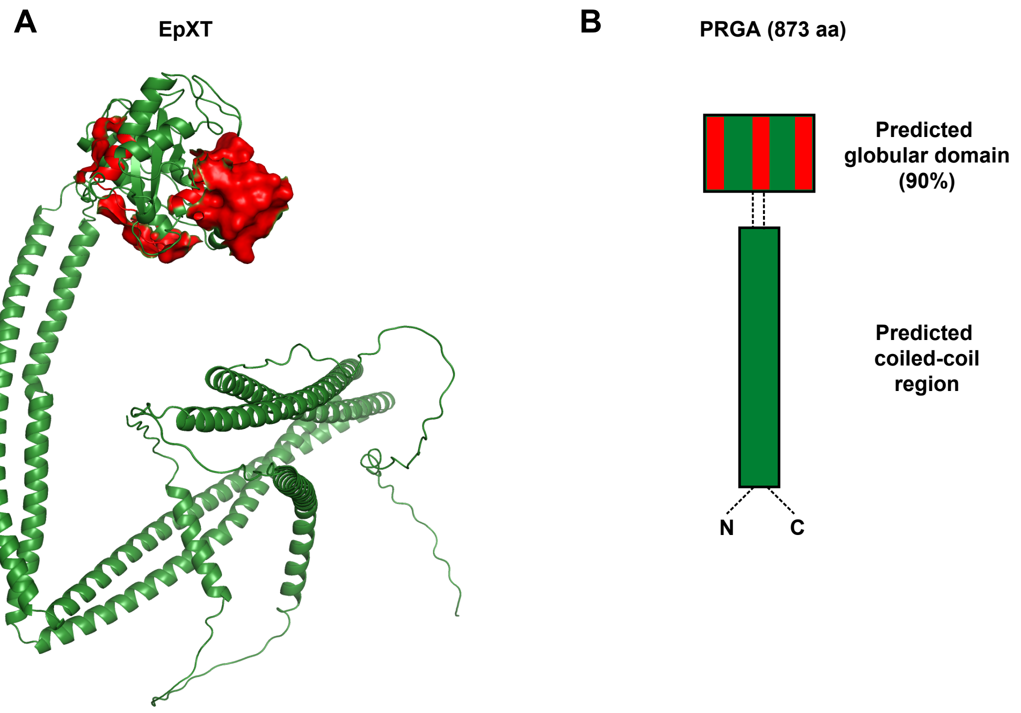
A
EpXT
B
PRGA (873 aa)
Predicted
globular domain
(90%)
Predicted
coiled-coil
region
N
C

A

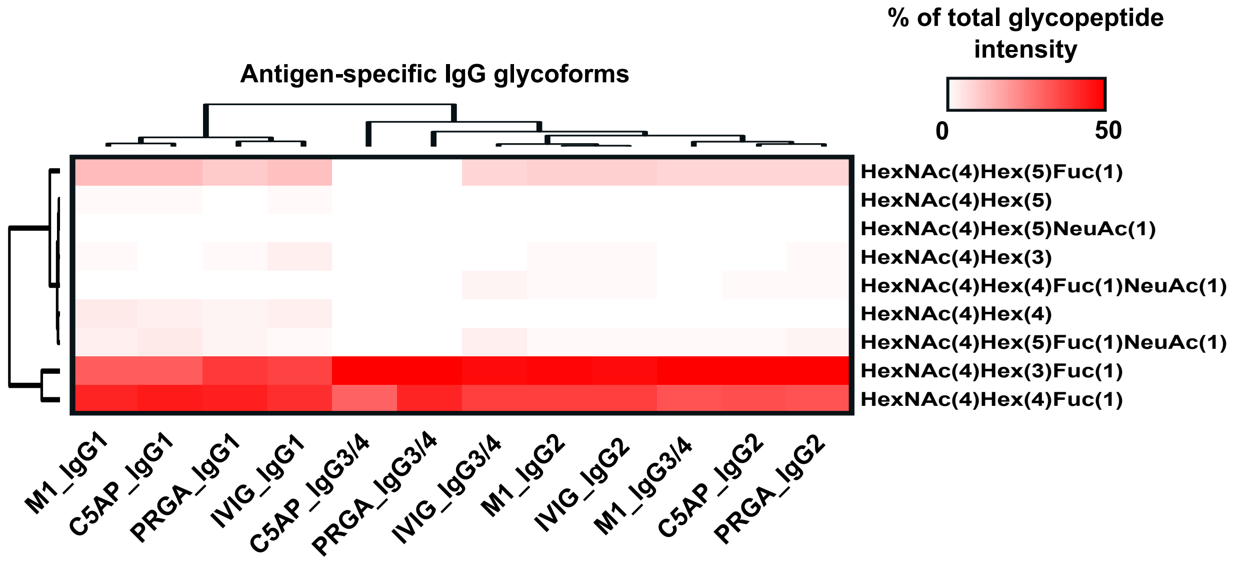

% of total glycopeptide intensity
Antigen-specific IgG glycoforms
0
50
HexNAc(4)Hex(5)Fuc(1)
HexNAc(4)Hex(5)
HexNAc(4)Hex(5)NeuAc(1)
HexNAc(4)Hex(3)
HexNAc(4)Hex(4)Fuc(1)NeuAc(1)
HexNAc(4)Hex(4)
HexNAc(4)Hex(5)Fuc(1)NeuAc(1)
HexNAc(4)Hex(3)Fuc(1)
HexNAc(4)Hex(4)Fuc(1)
M1_IgG1
C5AP_IgG1
PRGA_IgG1
IVIG_IgG1
C5AP_IgG3/4
PRGA_IgG3/4
IVIG_IgG3/4
M1_IgG2
IVIG_IgG2
M1_IgG3/4
C5AP_IgG2
PRGA_IgG2


B

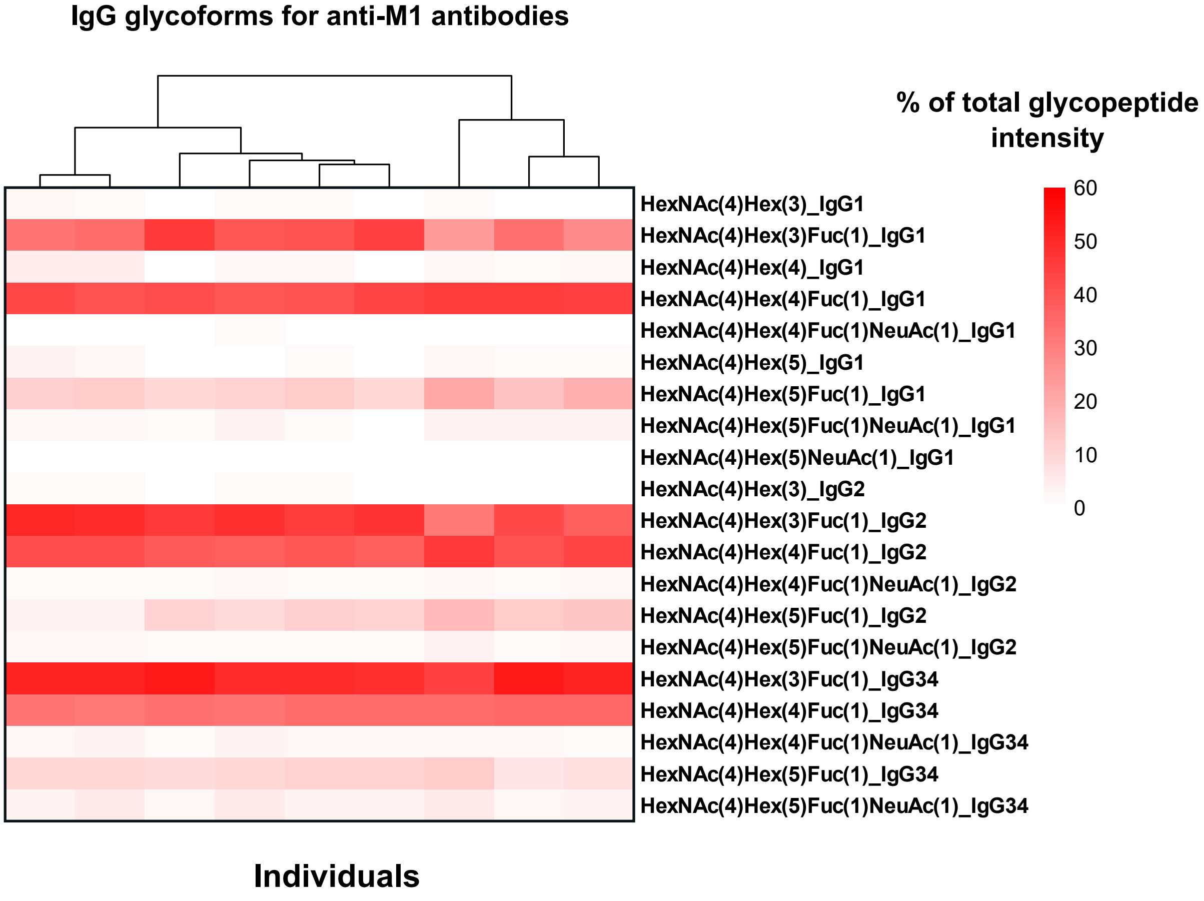

IgG glycoforms for anti-M1 antibodies
% of total glycopeptide intensity
60
50
40
30
20
10
0
HexNAc(4)Hex(3)_IgG1
HexNAc(4)Hex(3)Fuc(1)_IgG1
HexNAc(4)Hex(4)_IgG1
HexNAc(4)Hex(4)Fuc(1)_IgG1
HexNAc(4)Hex(4)Fuc(1)NeuAc(1)_IgG1
HexNAc(4)Hex(5)_IgG1
HexNAc(4)Hex(5)Fuc(1)_IgG1
HexNAc(4)Hex(5)Fuc(1)NeuAc(1)_IgG1
HexNAc(4)Hex(5)NeuAc(1)_IgG1
HexNAc(4)Hex(3)_IgG2
HexNAc(4)Hex(3)Fuc(1)_IgG2
HexNAc(4)Hex(4)Fuc(1)_IgG2
HexNAc(4)Hex(4)Fuc(1)NeuAc(1)_IgG2
HexNAc(4)Hex(5)Fuc(1)_IgG2
HexNAc(4)Hex(5)Fuc(1)NeuAc(1)_IgG2
HexNAc(4)Hex(3)Fuc(1)_IgG34
HexNAc(4)Hex(4)Fuc(1)_IgG34
HexNAc(4)Hex(4)Fuc(1)NeuAc(1)_IgG34
HexNAc(4)Hex(5)Fuc(1)_IgG34
HexNAc(4)Hex(5)Fuc(1)NeuAc(1)_IgG34
Individuals
